# *Burkholderia cenocepacia* epigenetic regulator M.BceJIV simultaneously engages two DNA recognition sequences for methylation

**DOI:** 10.1101/2024.01.20.576384

**Authors:** Richard Quintana-Feliciano, Jithesh Kottur, Mi Ni, Rikhia Ghosh, Leslie Salas-Estrada, Olga Rechkoblit, Marta Filizola, Gang Fang, Aneel K. Aggarwal

## Abstract

*Burkholderia cenocepacia* is an opportunistic and infective bacterium containing an orphan DNA methyltransferase (M.BceJIV) with roles in regulating gene expression and motility of the bacterium. M.BceJIV recognizes a GTWWAC motif (where W can be an adenine or a thymine) and methylates the N6 of the adenine at the fifth base position (GTWW**<u>A</u>**C). Here, we present a high-resolution crystal structure of M.BceJIV/DNA/sinefungin ternary complex and allied biochemical, computational, and thermodynamic analyses. Remarkably, the structure shows not one, but two DNA substrates bound to the M.BceJIV dimer, wherein each monomer contributes to the recognition of two recognition sequences. This unexpected mode of DNA binding and methylation has not been observed previously and sets a new precedent for a DNA methyltransferase. We also show that methylation at two recognition sequences occurs independently, and that GTWW**<u>A</u>**C motifs are enriched in intergenic regions of a strain of *B. cenocepacia’s* genome. We further computationally assess the interactions underlying the affinities of different ligands (SAM, SAH, and sinefungin) for M.BceJIV, as a step towards developing selective inhibitors for limiting *B. cenocepacia* infection.

## Introduction

*Burkholderia cenocepacia* is an opportunistic and infective bacterium, member of the *Burkholderia cepacia* complex (Bcc), that often leads to severe illness in immunocompromised patients ^1^. Antibiotics in clinical use against Bcc infection include the aminoglycoside Tobramycin and the β-lactams Aztreonam and Doripenem. *B. cenocepacia*’s ability to form biofilms and its antibiotic resistance highlight the importance of new approaches to counter its infectivity. Although bacterial DNA methyltransferases (MTases) are largely associated with restriction-modification (R-M) systems for defense against phages^2^, they can also play roles in gene regulation ^3–8^ and serve as potential drug targets^5,6,9^.

A DNA MTase was recently identified in *B. cenocepacia*, BCAM0992 (here referred to as M.BceJIV), deletion of which compromises the motility of the bacterium^10^. M.BceJIV is a 32 kDa MTase, part of the *trp* operon and potentially under the same regulation. It recognizes the GTWWAC motif (where W can be an adenine or a thymine) and methylates the N6 of the adenine at the fifth base position (GTWW**<u>A</u>**C)^10^. M.BceJIV is an orphan MTase, without a cognate restriction endonuclease, and is strongly conserved across the diverse species and strains of the *Burkholderia* genes, indicating its possible importance in the physiology of *Burkholderia* species^10^. Deletion of M.BceJIV leads to the upregulation of several genes with GTWWAC sites in their promoters, including the DNA helicase gene (rep) that has been shown to play a role in swimming motility in *Escherichia coli,* and may be correlated to the dysfunction in motility observed in M.BceJIV mutants^10^.

As an orphan MTase, M.BceJIV joins a growing list of bacterial MTases with roles outside of R-M and defense against foreign DNA^7^. Classical examples of orphan MTases are Dam in *Escherichia coli*, CcrM in *Caulobacter crescentus* and Dcm in *Escherichia* and *Salmonella* genra, functioning in DNA replication/repair, cell cycle regulation, and the expression of specific genes, respectively^7^. The recent discovery of an orphan MTase, CamA, in *Clostridioides difficile* with roles in the sporulation and motility of the bacterium highlights the therapeutic opportunities in targeting regulatory MTases in pathogenic bacteria^6,8^. We present here the crystal structure of a M.BceJIV/DNA complex that, unexpectedly, shows not one, but two DNA substrates bound to the M.BceJIV dimer. Remarkably, each monomer contributes to the recognition of two recognition sequences, a mode of DNA binding and methylation that has not been observed previously. As a key factor in *B. cenocepacia*’s motility, the M.BceJIV structure offers new opportunities in the design of therapeutics to control bacterial infection.

## Results

### Structure determination

M.BceJIV was cocrystallized with a pair of complementary 14-base-pair oligonucleotides with overhanging 5’-ends (A or T). The resulting DNA duplex contains a single M.BceJIV recognition site (GTTAAC). The cocrystals were obtained in the presence of sinefungin (in place of the natural cofactor SAM) and diffracted to ∼2.1 Å resolution with synchrotron radiation. The cocrystals belong to space group I4_1_ with unit cell dimensions of a = b = 137.7 Å, c = 167.3 Å, and α = β = γ = 90° and contain two M.BceJIV dimers, four DNA duplex substrates, and four sinefungin molecules in the crystallographic asymmetric unit (Table S1). The refined model consists of four M.BceJIV subunits (A, residues 30-279; B, residues 30-278; C, residues 30-279; D, residues 31-277), four DNA duplex substrates (DNA1, DNA2, DNA3, and DNA4), four sinefungin molecules, and 565 solvent molecules. M.BceJIV subunits A and B form a complex with DNA1 and DNA2 (complex 1), whereas subunits C and D form a complex with DNA3 and DNA4 (complex 2). The two complexes are very similar. The electron density for complex 1 is better defined than that for complex 2, and we describe the structure of complex 1 hereafter.

### Overall architecture

The M.BceJIV monomers A and B form a two-fold symmetric dimer that interacts extensively with two DNA duplexes (DNA1 and DNA2) containing the GTTAAC recognition motifs (Fig. 1). The target adenine (GTTA**<u>A</u>**C) flips out of each DNA duplex and enters the catalytic pockets of monomers A and B containing a sinefungin molecule (Fig. 1). Each M.BceJIV monomer contains the nine motifs (I-VIII and X) characteristic of amino MTases^11^ that together form an αβα “hub” consisting of a central eight-stranded β-sheet (β1-β8) with helices (αA, αB, αC, αF, αG and αH) on either side, from which the target recognition domain (TRD, composed of two short α-helices, αD and αE), protrudes outwards (Fig. 1). The TRD is a region of DNA MTases responsible for DNA recognition. Based on the linear order of the motifs, M.BceJIV belongs to the β class of amino MTases^11^, wherein the TRD (residues 167 to 194) is inserted between the N-terminal (IV-VIII) and the C-terminal (X and I-III) motifs. Motif I contains the FxGxG sequence for binding the cofactor SAM (or sinefungin in our case) and motif IV contains the (D/N)PP(Y/F) sequence for catalysis. The dimer interface between monomers A and B is extensive (∼1738 Å^2^), mediated mostly by helix αC, strands β4-β6, and loop-45 (between strands β4 and β5) from each monomer.

**Figure 1.**
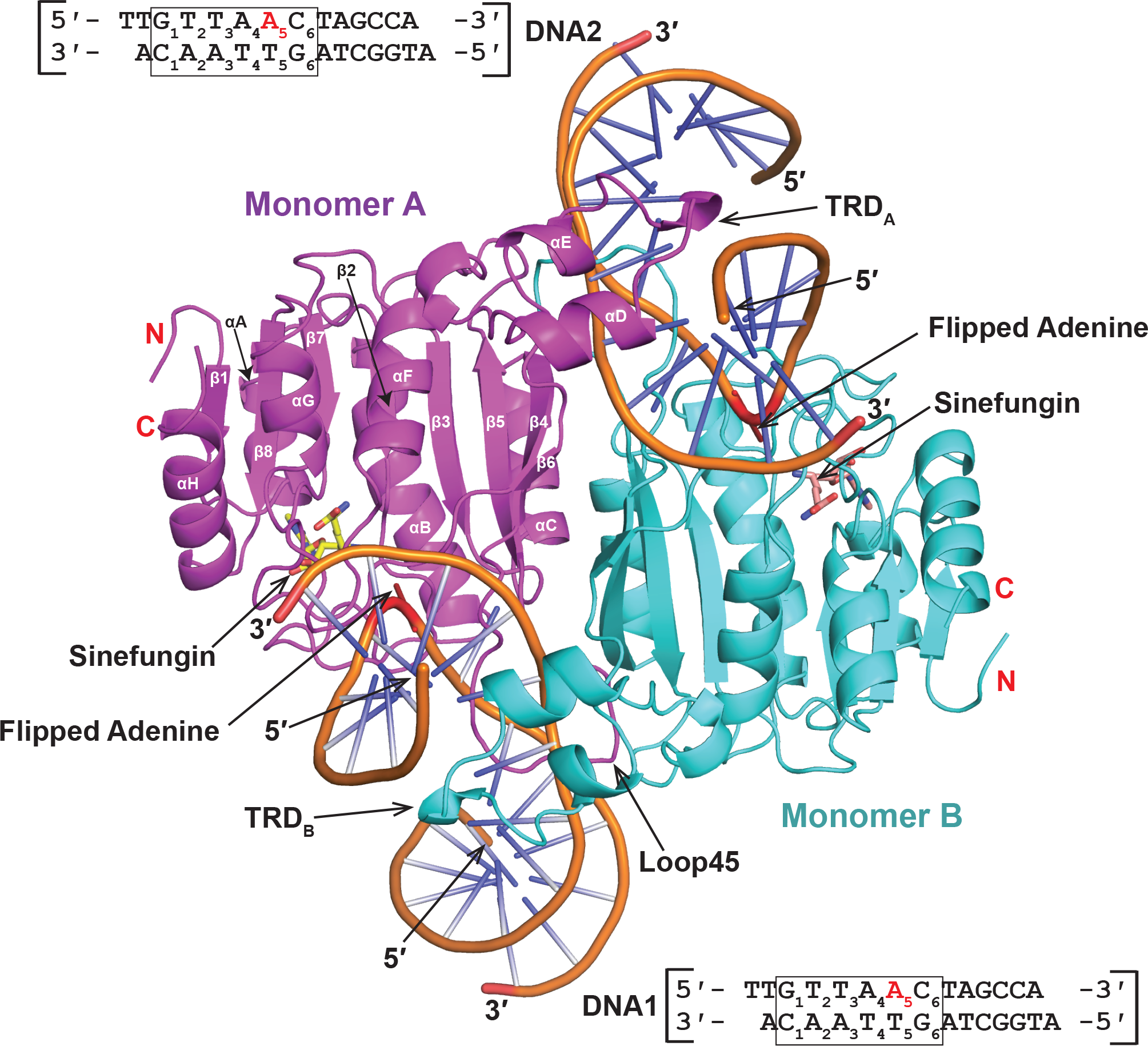
Overall structure of M.BceJIVΔ29-DNA-Sinefungin complex. M.BceJIV forms a homodimer complex with two DNAs and two Sinefungin molecules. Monomer A (magenta) receives the adenine to be methylated from DNA1 but its TRD reaches over to DNA2 and, conversely, monomer B (cyan) receives the adenine to be methylated from DNA2 but its TRD extends over to DNA1. Helices are labeled from αA to αH and strands are labeled as β1 to β8. Sinefungin is shown in a stick representation. The DNA substrate used for crystallization is shown and the recognition sequence is boxed and numbered.

Monomer A receives the adenine to be methylated from DNA1 but its TRD reaches over to DNA2 and, conversely, monomer B receives the adenine to be methylated from DNA2 but its TRD extends over to DNA1 (Fig. 1 and Fig. 2a,b). Thus, each monomer contributes to recognition of its own target DNA via its loop-45 and to the second DNA via its TRD (Fig. 2a,b). The TRD recognizes the DNA through the major groove, whereas the loop-45 recognizes the DNA through the minor groove. This rather unique mode of DNA recognition, whereby each M.BceJIV monomer contributes to the recognition of two DNA sequences is, to our knowledge, unprecedented for a DNA MTase.

**Figure 2.**
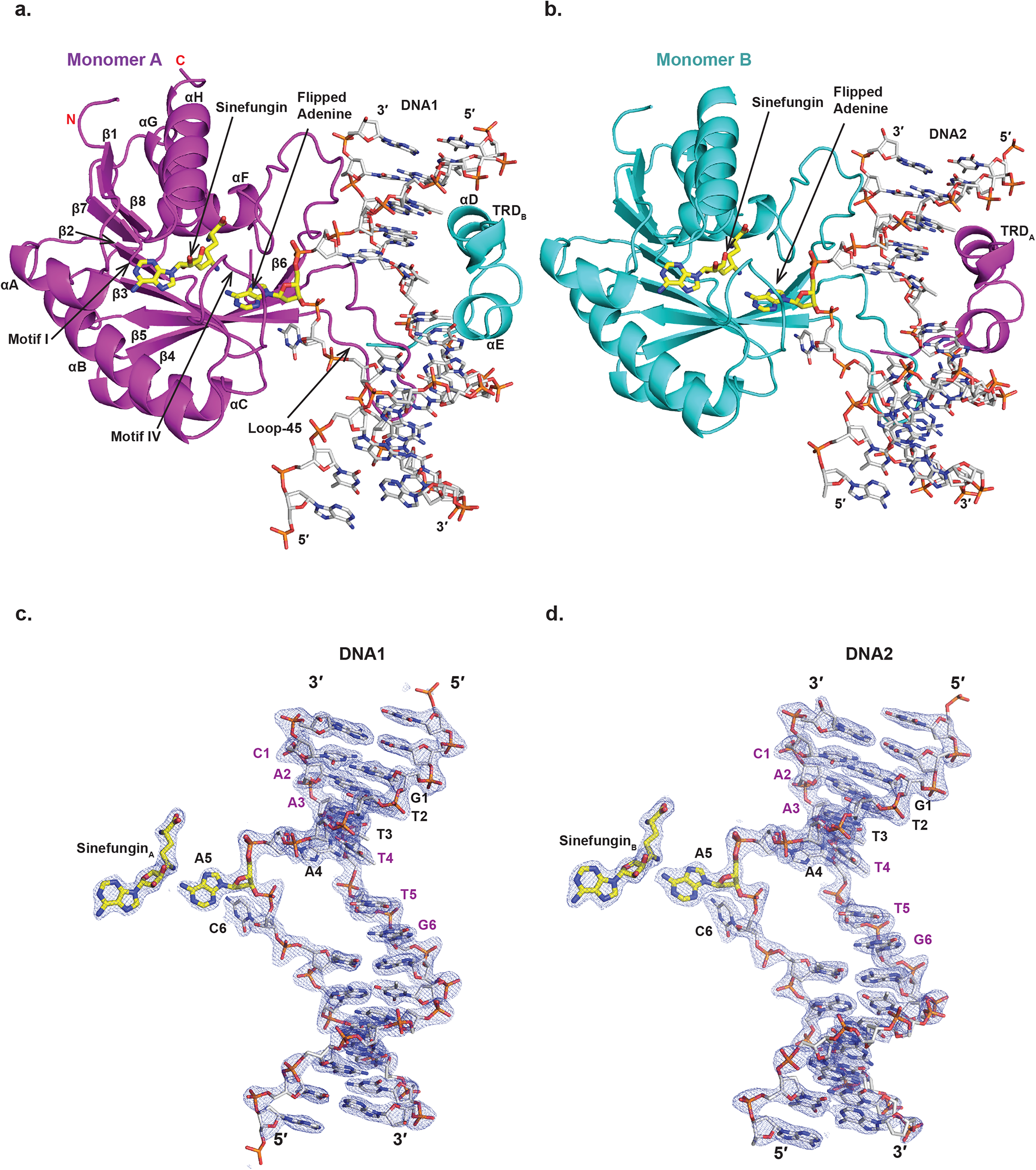
Structure of M.BceJIVΔ29 monomers bound to DNA and Sinefungin, and the electron density for DNA and Sinefungin molecules. Structure of monomer A bound to DNA1 and Sinefungin_A_ (**a**) and of monomer B bound to DNA2 and Sinefungin_B_ (**b**) showing DNA distortions and the TRDs. Each monomer contributes to recognition of its own methylation-target DNA via its loop-45 through the minor groove and to the recognition of a separate DNA via its TRD through the major groove. 2Fo-Fc map contoured at 1.5 σ is shown for DNA1 and Sinefungin_A_ (**c**) and for DNA2 and Sinefungin_B_ (**d**). The recognition sequence is labeled.

The DNAs (DNA1 and DNA2) are severely distorted from B-DNA conformation over the recognition sequence (GTTA<u>A</u>C) (Fig. 2c,d). In addition to the target adenine (GTTA**<u>A</u>**C, A5) being flipped out of the DNA helix, the cytosine 3’ to it (GTTA<u>A</u>**C, C6**) is also rotated out of the helix by ∼90 °, and the base pairs at the third and fourth positions (GT**TA**<u>A</u>C) are highly buckled at angles of ∼28 ° and 51°, respectively (Fig. 2). The major groove over the recognition sequence widens to ∼24 Å (from ∼19 Å in B-DNA) to accommodate the TRD and the minor groove widens to ∼18 Å (from ∼12 Å in B-DNA) to accommodate loop-45. Strikingly, residues 136 to 138 from loop-45 are lodged between the unpaired bases of the fifth and sixth base pairs (GTTA**<u>A</u>C**) and drive the adenine and the cytosine to their extrahelical positions (Fig. 3a).

**Figure 3.**
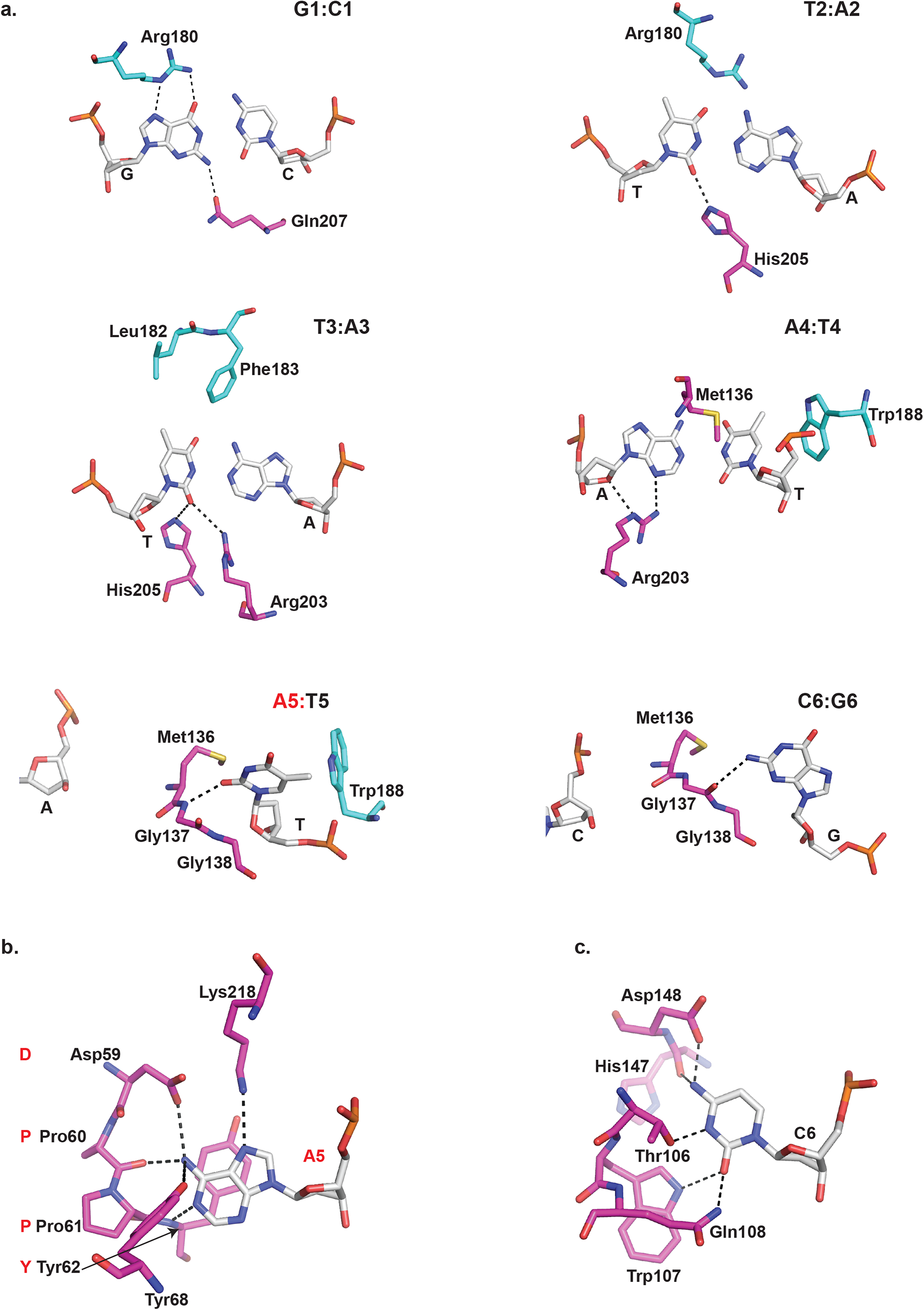
Molecular basis of target recognition. (**a)**, The first base pair (G1:C1) is specified by contacts with Gln207 of monomer A and Arg180 of TRD_B_; the second base pair (T2:A2) makes contact with His205 of monomer A and van der Waals contacts with Arg180 of TRD_B_; the third base pair (T3:A3) is specified by contacts with His205 and Arg203 of monomer A and van der Waals contacts with Leu182 and Phe183 of TRD_B_; the fourth base pair (A4:T4) makes contacts with Arg203 and van der Waals contacts with Met136 of monomer A (from loop-45) and van der Waals contacts with Trp188 of TRD_B_; at the fifth base pair (A5:T5), the thymine opposite the adenine to be methylated is specified by contacts with Gly137 of a short segment of monomer A (Met136-Gly137-Gly138 from loop-45), that embeds in place of the everted fifth adenine and sixth cytosines of the target strand, and van der Waals contacts with Trp188 of TRD_B_; at the sixth base pair (C6:G6), the guanine opposite to the everted cytosine makes contacts with Gly137 of monomer A (from loop-45). (**b)**, A close-up view of the flipped-out adenine that is accommodated in the catalytic cleft of monomer A and makes contacts with Asp59, Pro60 and Tyr62 of the conserved DPPY motif and with Tyr68 and Lys218 of monomer A. **c**, A close-up view of the flipped-out cytosine accommodated outside the catalytic cleft and makes contacts Asp148, His147, Thr106, Trp107, and Gln108 of monomer A. Monomer A is colored as magenta and the TRDB is colored as cyan.

### DNA recognition

For DNA1, the TRD of monomer B inserts into its major groove and the loop-45 of monomer A inserts into the minor groove. For DNA2 it is the converse, wherein TRD of monomer A inserts into its major groove and the loop-45 of monomer B inserts into the minor groove (Fig. 2a,b). Thus, each monomer recognizes both DNAs. Considering DNA1 (and the same applies to DNA2), the TRD contributes to the recognition of the first five of the six base-pairs of the recognition sequence (**GTTA<u>A</u>**C) (Fig. 3a). Specifically, Arg180 of the TRD makes bidentate hydrogen bonds with the O6 and N7 atoms of the outer guanine (**G**TTA<u>A</u>C) and van der Waals contacts with the C5 methyl group of the second thymine (G**T**TA<u>A</u>C), and Leu182 and Phe183 make van der Waals contacts with the C5 methyl of the third thymine (GT**T**A<u>A</u>C) (Fig. 3a). The fourth (A:**T**) and fifth (A:**T**) base pairs of the recognition sequence are specified by Trp188 that impinges on the C5 methyl groups of the thymines of these base-pairs (Fig. 3a). Loop-45 makes contacts through the minor groove, wherein Gln207 makes a hydrogen bond with the N2 amino group of the outer guanine (**G**TTA<u>A</u>C), His205 makes hydrogen bonds with the O2 atoms of the second and third thymines (G**TT**A<u>A</u>C), and Arg203 makes hydrogen bonds to the O2 atom of the third thymine (GT**T**A<u>A</u>C) and to the N3 atom of the fourth adenine (GTT**A**<u>A</u>C) (Fig. 3a). A short segment of loop-45 (Met136-Gly137-Gly138) embeds between the everted adenine and cytosine bases (GTTA**<u>A</u>C**) on the template strand and the partner thymine and guanine bases on the non-template strand (Fig. 3). Specifically, the main chain amide and carbonyl groups of Gly137 make direct hydrogen bonds with the thymine and guanine bases, respectively, facilitating their identity and acting as a wedge to evict the partner adenine and cytosine bases (GTTA**<u>A</u>C**) out of the DNA helix (Fig. 3). The flipped-out adenine of DNA1 enters the catalytic cleft of monomer A (where sinefungin is bound) and makes a series of specific contacts with residues Asp59, Pro60 and Tyr62 of the DPPY motif, conserved among amino MTases (described below). This adenine also makes additional polar interactions with Tyr68 and Lys218 (Fig. 3b). The flipped-out cytosine lies outside of catalytic cleft and its entire Watson-Crick face is engaged in hydrogen bonds with Thr106, Trp107, Gln108, His147, and Asp148 (Fig. 3c). In addition, Met136 of Loop-45 impinges on the A:T base pair at the fourth position (GTT**A**<u>A</u>C) and contributes to the highly buckled state (∼51°) of this base pair (Fig. 3a, A4:T4). Overall, M.BceJIV recognizes DNA through a combination of base-specific contacts and distortion of the DNA. That is, the free energy to adopt the distorted configuration of the DNA with evicted and buckled bases (Fig. 2c,d) is favored by the excess of A:T base pairs within the recognition sequence (GTWW**<u>A</u>**C).

### DNA methylation

As noted above, the target adenine of DNA1 inserts into the catalytic cleft of molecule A and interacts with the conserved DPPY motif. Conversely, the target adenine of DNA2 enters the catalytic cleft of molecule B. The PPY sequence of the DPPY motif in molecules A and B runs along the Watson-Crick edge of the extrahelical adenine rather than the Hoogsteen edge as in γ-class of amino MTases^12^. Considering DNA1 (and the same applies to DNA2), the extrahelical adenine stacks between Tyr62 and Thr216 and its N6 amino group makes hydrogen bonds with the Pro60 main chain carbonyl O and the Asp59 side chain OD1 atoms and a polar interaction with the side chain OH of Tyr68, and its N7 atom makes a hydrogen bond with Lys218. There are also putative weaker out of plane hydrogen bonds between its N1 atom and the main chain amide NH of Tyr62 and the side chain OH of Tyr68. Overall, this pattern of hydrogen bonding specifies an adenine in the catalytic cleft and positions its N6 amino group close to the methyl group of modeled SAM for methyl transfer by an in-line nucleophilic attack. To avoid catalysis and methyl transfer in our complex, we crystallized M.BceJIV in the presence of a SAM analog, sinefungin, which has a nontransferable amino group in place of the methyl group. Sinefungin is bound towards the C-terminal ends of strands β7 and β8 with the adenine ring stacked between Phe241 and Ile263, the sugar hydroxyls making hydrogen bonds with Glu262 OE1 and OE2 atoms, and the carboxyl end making hydrogen bonds with the main chain and side chain atoms of Ser244, main chain NH of Lys218 and OG1 atom of Thr246 (Fig. 4). The non-transferable amino group of sinefungin is located only 2.9 Å from the N6 atom of the extrahelical target adenine and is involved in polar interactions with the hydroxyl group of Tyr68 and with both the main chain and side chain atoms of Asp59.

**Figure 4.**
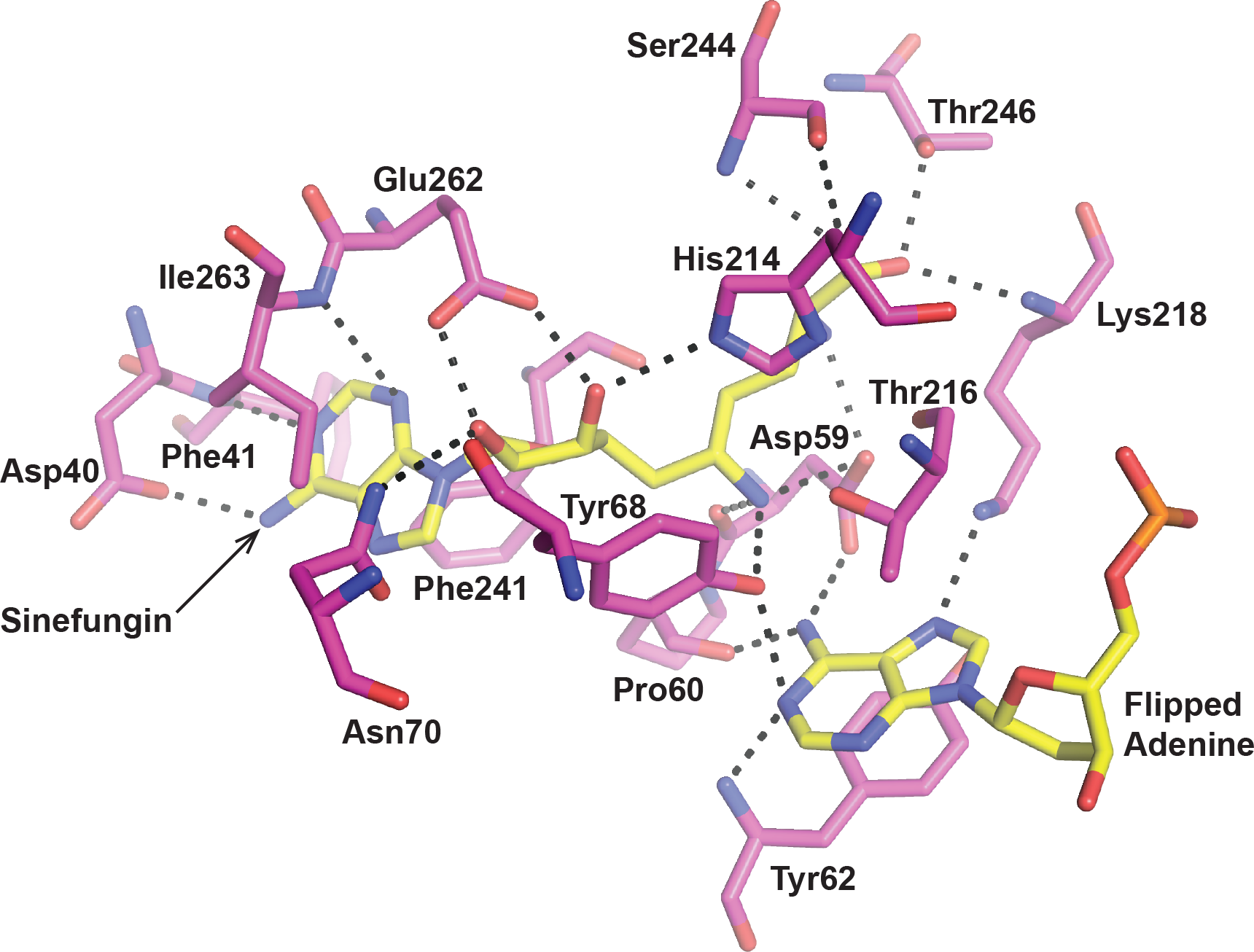
Interactions of M.BceJIVΔ29 with Sinefungin and the flipped adenine. A close-up view of specific M.BceJIVΔ29-Sinefungin interactions and residues shared between Sinefungin and the flipped-out adenine. Sinefungin is tightly bound within the MTase domain via extensive hydrogen bonding (dashed lines) and hydrophobic contacts.

### M.BceJIV methylates the two DNA sites independently

The finding that M.BceJIV can simultaneously bind two DNA substrates prompts the question of whether the binding and methylation of one DNA substrate facilitates the binding and methylation of the second DNA substrate. Many restriction enzymes are found to require two or more DNA sites for cleavage^13–16^ but whether a similar cooperation exists for a DNA MTase is unclear. To determine if methylation on a motif is dependent on the methylation of another motif, a set of experiments coupling MTase activity of full-length M.BceJIV with the endonuclease activity of HpaI (NEB) were performed. HpaI is a restriction endonuclease that introduces a double-stranded break in the middle of the sequence 5’-GTTA**<u>A</u>**C-3’, which is a motif that contains an adenine (underlined) that gets methylated by M.BceJIV. We supplied the second motif *in trans* or *cis* and asked whether it leads to stimulation in methylation of the first site (see Supplementary Fig. S1). For these experiments, we constructed a plasmid carrying a single GTTAAC sequence that serves as recognition site for both M.BceJIV (GTWW**<u>A</u>**C) and the restriction endonuclease HpaI (GTT**^▾^**AAC). The addition of HpaI to the linearized plasmid led to complete cleavage that could be visualized on an agarose gel with ethidium bromide. As expected, the addition of M.BceJIV and cofactor SAM to the mixture led to inhibition of cleavage by HpaI. At appropriate concentrations of the enzymes and times for the reaction corresponding to ∼50% inhibition of DNA cleavage, we tested whether the addition of an oligonucleotide *in trans* carrying the recognition sequence for M.BceJIV (GTAA**<u>A</u>**C), but not for HpaI, would stimulate methylation of the first site on the linearized plasmid (monitored as reduction in cleavage by HpaI). The assay is analogous to that reported previously for the MmeI family of restriction enzymes which revealed stimulation of cleavage at one site when another is supplied *in trans*^17^. As shown in Fig. 5a, we observed no stimulation in methylation by M.BceJIV of the site on the plasmid when another was provided *in trans*. If anything, there is a slight inhibition in methylation of the first site when the second was provided *in trans*, presumably due to titration of the enzyme by the second site (Fig. 5a). Thus, even though the M.BceJIV dimer, from our structure, can simultaneously bind and methylate two recognition sequences, it appears to do so independently without cooperativity between the sites. To test whether this was also the case when the two sites are *in cis*, we constructed a plasmid carrying two recognition sequences, one for M.BceJIV/HpaI (site 1, GTT**^▾^**A**<u>A</u>**C) and another for just M.BceJIV (site 2, GTAA**<u>A</u>**C) (Supplementary Fig. S1) and monitored for HpaI cleavage of site 1 as a proxy for the degree of methylation. As shown in Fig. 5b, there is no stimulation of methylation of site 1 when the plasmid contained another M.BceJIV recognition sequence *in cis*. Thus, the M.BceJIV dimer can simultaneously bind and methylate two recognition sequences but this occurs independently and without cooperativity between the sites.

**Figure 5.**
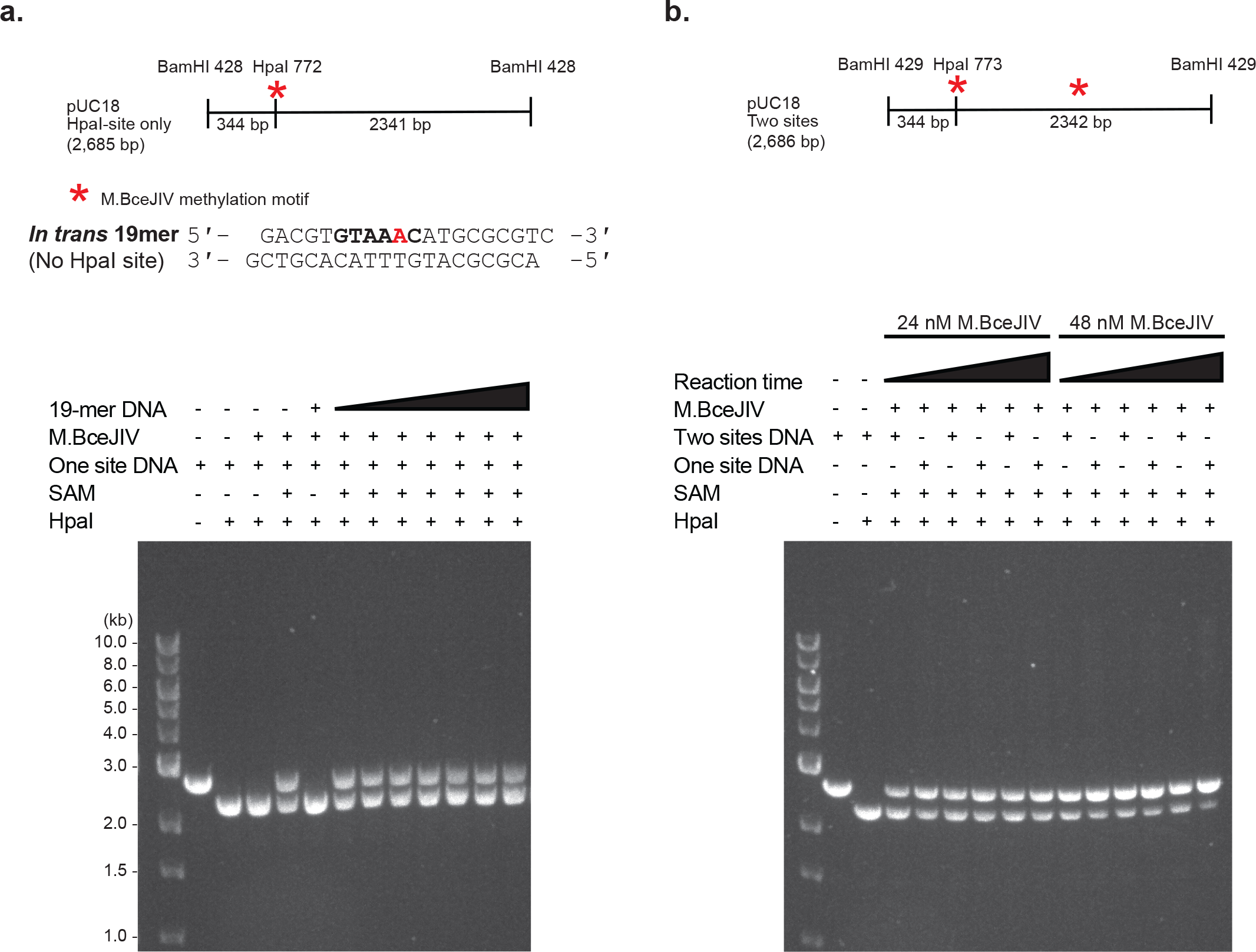
Addition of a second DNA methylation site *in trans* or *in cis* does not assist the methylation of the first DNA site. (a) *in trans* experiment, HpaI digestion of 24 nM BamHI-linearized pUC18 containing an HpaI site only with increasing concentrations (0 nM to 50 nM) of a 19 mer DNA duplex *in trans* containing a M.BceJIV methylation motif and no HpaI site in the presence of 24 nM M.BceJIV and 20 μM SAM in a 50 s reaction. (b) *in cis* experiment, HpaI digestion of 24 nM BamHI-linearized pUC18 containing an HpaI site only (or both an HpaI site and another M.BceJIV methylation site *in cis*) with increasing reaction time (1 min to 5 min) in the presence of 24 nM or 48 nM M.BceJIV and 20 μM SAM.

### Genomic distribution of M.BceJIV recognition sites

To understand the genome-wide distribution of the M.BceJIV recognition sites (5’-GTWWAC-3’), we conducted a genomic analysis of two *B. cenocepacia* strains, J2315 and K56-2, each with an approximate size of 8 Mb. Both strains comprise three chromosomes and one plasmid (Fig. 6a,b). Within strain J2315, we identified a total of 982 GTWWAC sites with a wide range of distances between each two neighboring sites (Fig. 6c). While most neighboring sites were about 7 kb from each other, some sites are as close as <10 bp and as far as >10 kb from their neighboring sites. The other strain K56-2, with 933 GTWWAC sites, exhibited a comparable pattern in the distribution of GTWWAC sites and distances between neighboring sites as strain J2315 (Fig. 6d). When we examined the relative distribution of GTWWAC sites between intergenic and intragenic regions, we found that 35.7% of the motif sites are in the intergenic regions of the J2315 genome, representing nearly 3-fold enrichment considering the size of intergenic regions in the genome (∼12.5%). Hypothesizing that this enrichment in the intergenic region is likely due to the unusual AT-rich 4-mer (TWWA) in the GTWWAC motif, we performed a Markov chain motif frequency analysis^8^ and found that the observed enrichment of GTWWAC in the intergenic region is indeed largely driven by the AT rich nature of intergenic regions compared to intragenic regions (Fig. 6e,f). These motif distribution analyses highlight that the GTWWAC methylation motif may mediate the competition between M.BceJIV and other DNA binding proteins such as transcription factors to facilitate epigenetic regulation^3^.

**Figure 6.**
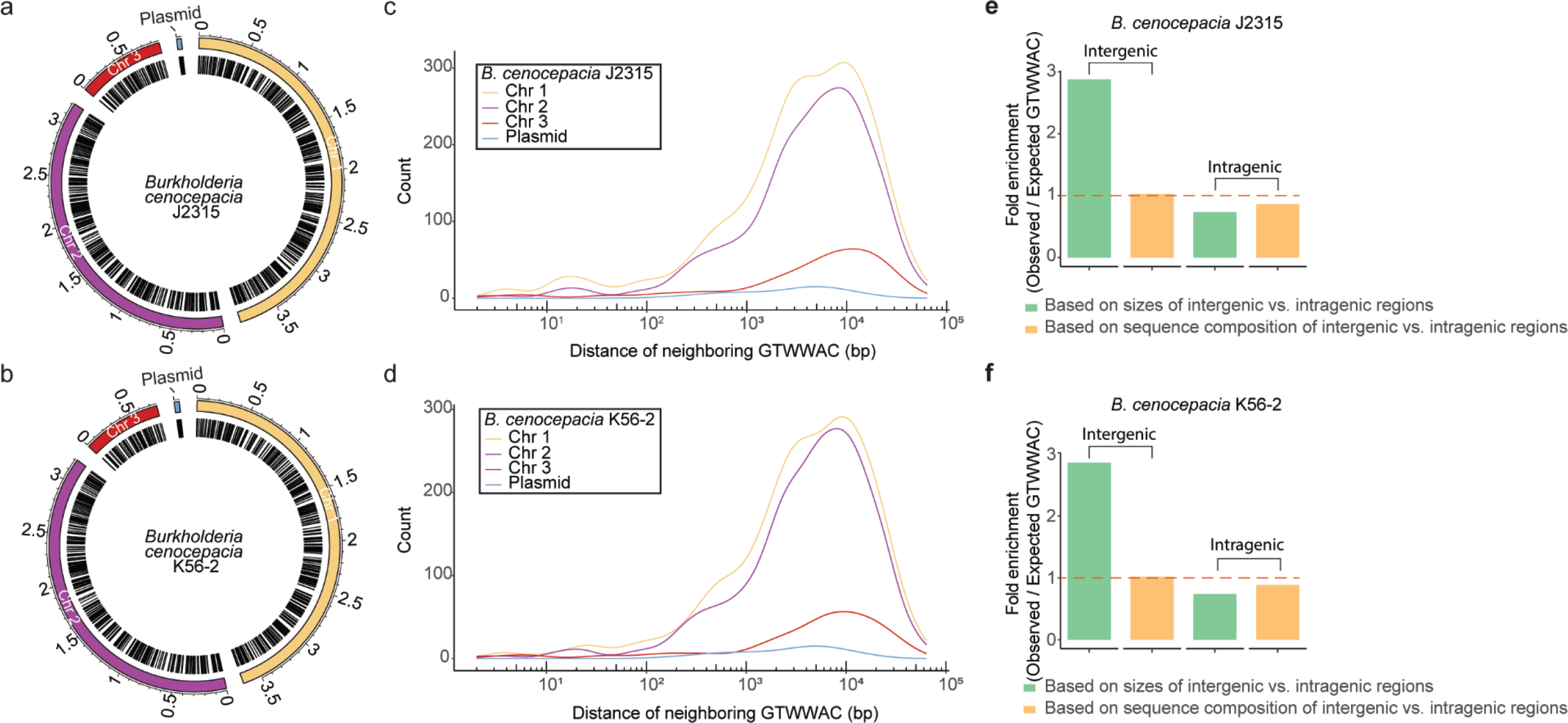
Distribution of GTWWAC motif sites across two strains of *B. cenocepacia*. (**a & b),** Circos plot showing the genome-wide distribution of GTWWAC sites along the three chromosomes and a plasmid of *B. cenocepacia* J2315 and *B. cenocepacia* K56-2, respectively. (**c & d),** Density plot for the distribution of distances between each two neighboring GTWWAC sites in *B. cenocepacia* strains J2315 and K56-2. (**e & f**), Enrichment of GTWWAC in the intergenic regions (observed frequency vs. expected frequency) in the two strains.

Given that the structures of numerous DNA MTases are presently available, we searched the PDB database for MTases with high similarity. The enzyme RsrI from *Rhodobacter sphaeroides* displayed the highest degree of similarity, sharing 40.0% amino acid identity with the strains studied^18^. Additionally, an amino acid identity of 36.2% was observed between M.BceJIV and MboIIA from *Moraxella bovis*, the latter being an enzyme that methylates the 3’ adenine of GAAGA and usually forms a dimer when it is not bound to DNA^19^.

### Binding and kinetic parameters

We measured the binding affinities of SAM, SAH and sinefungin for M.BceJIV by isothermal titration calorimetry (ITC). From the ITC analysis, SAM and sinefungin bind M.BceJIV with similar affinities (*K*_D_s of ∼ 8 μM) but SAH binds with approximately a two-fold weaker affinity (*K*_D_ of ∼17 μM) (Fig. 7a). Energy-minimized docking structures of SAM and SAH at the crystallographic binding pockets of sinefungin in the M.BceJIV-DNA dimeric complex surrounded by an explicit water environment reveal differences in the interactions between the three ligands and the protein, particularly in regions where their chemical structures vary (Fig. 7b). Sinefungin is stabilized in part by an electrostatic interaction that its non-transferable, positively charged amino group establishes with Asp59, as well as hydrogen bonds formed with the main chain carbonyl oxygen atom, one of the side chain carboxylate oxygen atoms of Asp59, and the hydroxyl group of Tyr68. Similarly, SAM is stabilized by an electrostatic interaction between its positively charged sulfonium group and Asp59, along with possible unconventional C-H··O hydrogen bonds between its sulfonium methyl group and the main chain carbonyl oxygen atom, one of the side chain carboxylate oxygen atoms of Asp59, and the hydroxyl group of Tyr68. Additionally, there may be possible sulfonium−π interactions with Tyr68. Notably, C-H··O bonds have been postulated in SAM-dependent methyltransferases^20^ due to the polarization of the methyl carbon within the positively charged sulfonium group. A recent study has demonstrated that this polarization also contributes to making sulfonium−π interactions stronger than sulfur−π and ammonium-π interactions, mostly because of the additional methyl−π interaction offered by the sulfonium group^21^. Unlike sinefungin and SAM, SAH is unable to participate in equivalent electrostatic, polar, and methyl-π interactions, which may account for its lower affinity.

**Figure 7.**
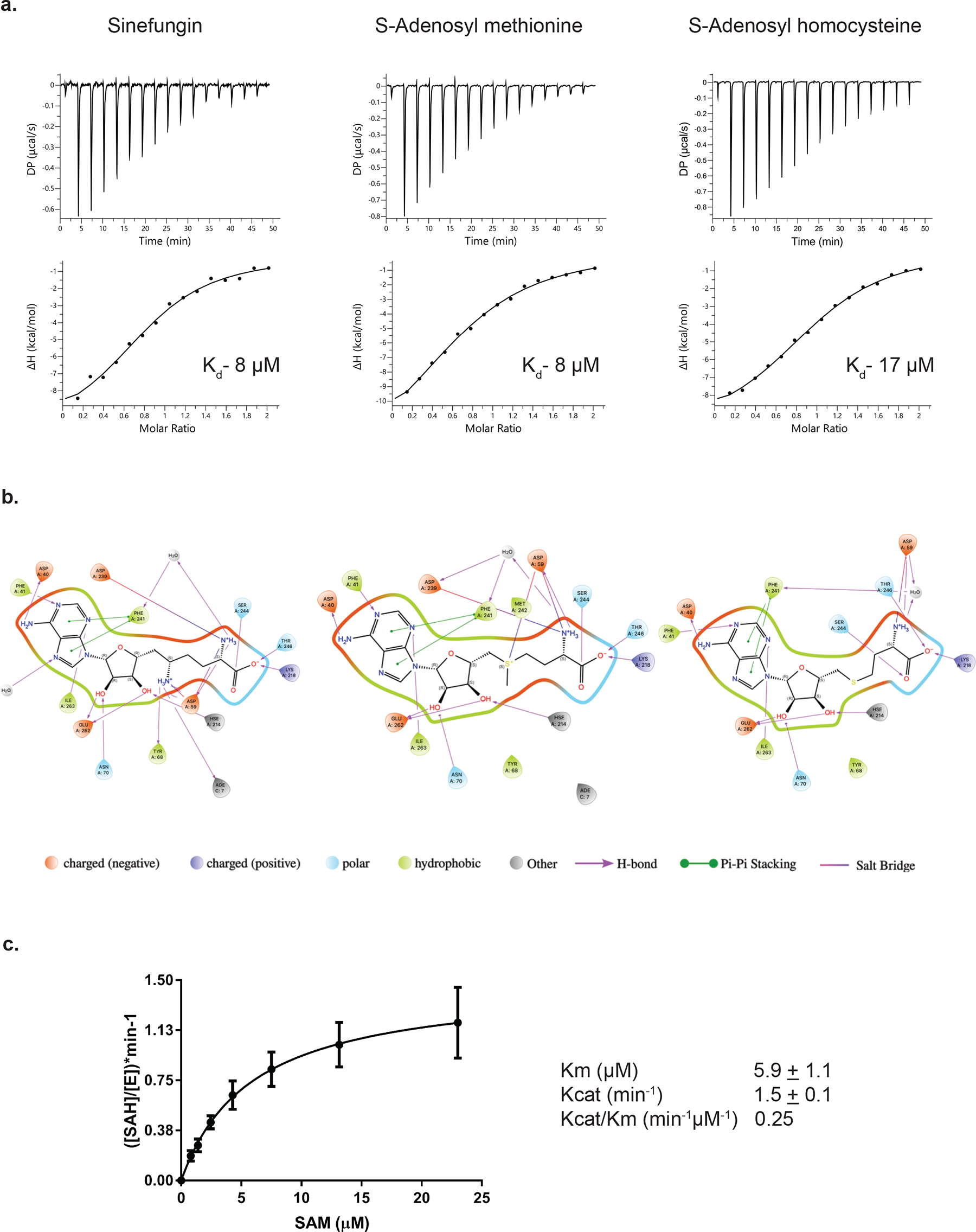
ITC analysis of M.BceJIVΔ29 with SAM, SAH and Sinefungin, and kinetics of M.BceJIV using a bioluminescence assay. (**a)**, ITC titration data for Sinefungin (left), SAM (middle) and SAH (right) with M.BceJIV**Δ**29. The equilibrium dissociation constants (K_D_) were derived from the resulting binding isotherms. (**b)**, Two-dimensional protein-ligand interaction diagrams generated using energy-minimized conformations of the crystal structure of M.BceJIVΔ29 bound to Sinefungin (left) and predicted structures of SAM and SAH in M.BceJIVΔ29 (middle and right, respectively) embedded in an explicit water environment. The Ligand Interaction script in Maestro (Schrödinger Inc., www.schrodinger.com) was used to generate these diagrams with default parameters. Among the protein residues within a 4 Å distance of the ligands, only those that interact with the ligands according to Maestro’s default parameters (see details in Table S2) are displayed. (**c)**, Kinetics of SAH byproduct formation by varying concentrations of the methyl donor SAM. Summary of kinetic parameters are also shown.

We next measured the M.BceJIV kinetic parameters by measuring concentrations of the reaction product SAH under varying concentrations of SAM. From the kinetic analysis, M.BceJIV has *K*_m_ and *k*_cat_ values of ∼5.9 μM and ∼1.5 min^-1^, respectively (Fig. 7c). The *K*_m_ value is approximately similar to the *K*_D_ value for SAM determined from ITC analysis. The *k*_cat_ value is at the lower end of values reported for other orphan DNA MTases, namely *E. coli* Dam (∼0.14 min^-1^)^22^, *C. difficile* CamA (∼5.4 min^-1^)^5^, and *C. crescentus* CcrM (∼5.2 min^-1^)^23^. Like M.BceJIV, CcrM is also a dimer and belongs to the β class of amino MTases, but unlike M.BceJIV, it engages only a single DNA substate as a dimer^24^. Thus, even though the M.BceJIV dimer engages two DNA substrates for methylation, it does not lead to a faster catalytic turnover of the substrate than CcrM.

## Discussion

We present here a high-resolution crystal structure of the M.BceJIV/DNA/sinefungin ternary complex. M.BceJIV is only the third β class of amino MTases to be visualized with DNA. The first one, EcoP15I, revealed dimerization as the defining feature of this class of MTases and uncovered a remarkable division of labor, where one MTase subunit recognizes the DNA, while the other MTase subunit methylates the target adenine^25^. The second, cell-cycle regulated CcrM, also revealed a dimer, wherein the two subunits cooperate in pulling apart the DNA strands over the recognition sequence^24^. In both the EcoP15I and CcrM complexes, the MTase dimer binds a single DNA duplex. M.BceJIV is also a dimer but it engages not one, but two DNA duplexes, wherein each subunit contributes to the binding and recognition of both DNA duplexes. Thus, even though M.BceJIV, EcoP15I and CcrM belong to the same β class of amino MTases and are homodimers, there is surprising diversity in how they distribute labor between the subunits for DNA binding and methylation.

The ability of M.BceJIV to engage two recognition sequences posits an ability to co-methylate two DNA sites at the same time, whether they occur *in trans* or *cis*. The *cis* sites on a bacterial chromosome could be potentially far apart with the DNA in between them looped out when they bind M.BceJIV dimer. To test whether co-methylation is cooperative, we monitored the extent of methylation at one DNA site when another is supplied *in trans* or *cis*. From these experiments, the methylation on one site does not stimulate the methylation of the second site, suggesting that co-methylation on two DNA sites occurs independently without cooperativity. From our bioinformatic analysis of the *B. cenocepacia* genome, the M.BceJIV recognition sequences are distributed with a wide range of distance between each two neighboring sites. While most neighboring sites were about 7 kb from each other, some sites are as close as a few base pairs and as far as >10 kb from their neighboring sites. Intriguingly, the sites are significantly enriched in intergenic regions, a reflection perhaps of the regulatory role of M.BceJIV in gene transcription.

Bacterial DNA MTases are commonly part of the R-M systems for defense against phages^2^. As an orphan MTase (unassociated with a restriction enzyme), M.BceJIV joins a growing number of bacterial MTases with putative roles outside of phage defense^4,7^. A recent study in *B. cenocepacia* strains J2351 and K56-2 shows a role for M.BceJIV in the epigenetic regulation of gene expression related to cellular motility, making M.BceJIV a viable therapeutic target for limiting *B. cenocepacia* infections^10^. A potential set of therapeutics is small molecules that compete with binding of the cofactor SAM to M.BceJIV. Indeed, many SAM-mimetic and non-SAM mimetic compounds have been developed to inhibit protein MTases in the field of cancer^26,27^ and it would be meaningful to test some of these for inhibition of M.BceJIV as starting lead compounds. The structure of M.BceJIV provides a basis for optimizing these lead compounds for increased potency and selectivity through iterative cycles of crystallography, chemical synthesis, and assays for *B. cenocepacia’s* motility. Interestingly, *B. cenocepacia* wild-type cells treated with sinefungin are less motile (to a similar extent as a M.BceJIV deletion mutant), consistent with the inhibition of M.BceJIV in a cellular context^10^. One issue with sinefungin as a potential therapeutic is that it is a pan MTase inhibitor with potential off-target effects, including the inhibition of host MTases in any treatment regimen. However, it is noteworthy from our studies that sinefungin binds M.BceJIV with a similar affinity as SAM (*K*Ds of ∼ 8 μM) due to favorable interactions between its non-tranferable positively charged amino group and Asp59 and Tyr68; positing sinefungin as a good starting scaffold for the design of SAM-mimetics selective for M.BceJIV for inhibition of *B. cenocepacia*’s motility. Altogether, inhibition of bacterial DNA MTases for controlling bacterial infections has gained traction over the past few years^5,6,8^. A notable example is the development of selective and potent inhibitors (*K*_D_ of ∼0.2 μM) for the DNA adenine MTase CamA for controlling *C. difficile’s* sporulation and biofilm formation^9^.

In summary, we present here a high-resolution crystal structure of the M.BceJIV/DNA/sinefungin ternary complex. The structure reveals a mode of DNA recognition and methylation not observed previously with other DNA MTases, wherein each M.BceJIV monomer contributes to the recognition of two DNA sequences. Given the role of M.BceJIV in regulating *B. cenocepacia’*s cellular motility, it is a viable therapeutic target, and the structure presented here will be invaluable in developing selective and potent inhibitors of M.BceJIV for limiting *B. cenocepacia* infections.

## Methods

### Expression and purification

For the expression of full-length N-terminal His tagged M.BceJIV (Uniprot Accession number B4EFK1), the codon optimized gene was cloned into a pET28-smt3 vector. For crystallization, a construct of M.BceJIV without the first 29 N-terminal residues (M.BceJIV**Δ**29) was cloned in the pET28-smt3 vector with N-terminal 6xHis tag. The plasmids were transformed into *E. coli* NiCo21(DE3) cells (NEB) and the cells were grown in LB media supplemented with 50 µg/mL Kanamycin at 37°C until OD_600_ reached 0.8. Protein expression was induced with 0.6 mM isopropyl β-D-1-thiogalactopyranoside and cultures were incubated for 17 h at 18 °C. The cell pellets from 4 L were resuspended in 150 mL binding buffer (50 mM Potassium phosphate pH 8.0, 500 mM Sodium chloride (NaCl), 5% glycerol, 0.01% triton X-100, 5 mM β-mercaptoethanol, 30 mM Imidazole and 10 μM Zinc chloride) containing 1 mM Phenylmethylsulfonyl fluoride and protease inhibitors. Cells were sonicated in ice for 10 min with 15 s ON and 2 min OFF and cell debris were removed by centrifugation at 18,000 rpm. The supernatant was filtered and incubated with 5 mL of Ni-NTA beads (Thermo Scientific) for 1 h. The beads were then washed with 100 mL binding buffer followed by 100 mL high salt buffer (binding buffer with additional 500 mM NaCl) and 30 mL binding buffer. The protein was then eluted with elution buffer (binding buffer with 500 mM imidazole) and the His tag was cleaved using Ulp1-protease by dialyzing overnight at 4 °C against 1 L binding buffer with 100 mM NaCl. To remove the cleaved His-tag, the protein was passed through a 5 mL HisTrap HP column (GE Healthcare), and the cleaved protein was collected in the flow through. The protein was further purified using a 5 mL HiTrap Heparin HP column (GE Healthcare), and then eluted with a linear gradient of 0.1-0.5 M NaCl. The peak elution fractions were pooled and concentrated to 2 mL using a Amicon Ultra 10K MWCO centrifugal filter at 4 °C and was loaded into a HiLoad Superdex 200 16/60 column (GE healthcare) equilibrated with 20 mM HEPES pH 8.0, 300 mM NaCl, 5% glycerol and 0.5 mM tris(2-carboxyethyl)phosphine. The purified protein was concentrated to >35 mg/mL, aliquoted and stored at -80 °C.

### Crystallization and structure determination

For crystallization experiments, the ternary complex was reconstituted by mixing 20 mg/mL M.BceJIV**Δ**29, 2.5 mM Sinefungin and 0.83 mM DNA duplex (5′-TTGTTA**A**CTAGCCA-3’ and 3′-ACAATTGATCGGTA-5′) in size exclusion buffer. The mix was incubated for at least 1 h in ice followed by centrifugation at 13,000 rpm at 4 °C for 10 min to remove any precipitate. Initial crystallization screening was done with commercially available screens using an Oryx Nano (Douglas Instruments) in a sitting-drop format with 0.25:0.25 µL protein-reservoir ratio at 17 °C. The initial crystals were obtained within 2 d in solution containing 0.10% w/v n-Octyl-β-D-glucoside, 0.1 M Sodium citrate tribasic dihydrate pH 5.5, and 22% w/v Polyethylene glycol 3350. The crystals were further optimized by varying the pH of the crystallization buffer, NaCl concentration in the sample mix and percent of precipitant in a hanging-drop format using 1:1 and 1:1.25 µL protein-reservoir ratios. Crystals were cryoprotected in a stepwise manner with reservoir solution containing 5-25% sucrose and flash-frozen in liquid nitrogen after 1-5 min incubation.

X-ray diffraction data were collected under cryogenic conditions at the National Synchrotron Light Source II 17-ID-2 beamlines at the Brookhaven National Laboratory. The diffraction data were processed using autoPROC and STARANISO (Global Phasing Ltd.). The structure was solved by molecular replacement using Phaser-MR with the AlphaFold^28^ model of M.BceJIV as the search model and to model the DNA molecules the PDB entry 6PBD was utilized as a reference. Subsequent iterative manual binding was performed with Coot^29^ and refinement with Phenix Refine^30^. A ligand restraint file for Sinefungin was generated with eLBOW^31^ from the PHENIX suite. PyMOL (Schrödinger LLC) was used to prepare all the structure figures.

### Methylation-endonuclease coupled assays

For methylation-endonuclease coupled assays, pUC18 plasmids were linearized by digestion with BamHI prior to these experiments. For the *in trans* experiments, increasing concentrations (0 nM to 50 nM) of a 19 mer DNA duplex (5’-GACGT<u>GTAA**A**C</u>ATGCGCGTC-3’ and 3’-GCTGCACATTTGTACGCGCA-5’) containing a M.BceJIV methylation motif and no HpaI endonuclease site was added *in trans* when indicated and the mix was further incubated for 30 min at room temperature in reaction buffer (20 mM Tris pH 8.0, 50 mM NaCl, 1 mM ethylenediaminetetraacetic acid, 3 mM magnesium chloride, 1 mM dithiothreitol and 0.1 mg/ml Bis(trimethylsilyl)acetamide). For the *in cis* experiments, methylation-endonuclease activity was assayed in 20 µL reaction mixes by incubating 24 nM or 48 nM of full-length M.BceJIV with 24 nM of pUC18 plasmids containing an HpaI site only (or both an HpaI site and another M.BceJIV recognition site *in cis*) for 30 min at room temperature in reaction buffer. For all the experiments, the methyltransferase reactions were started by adding 20 μM SAM and proceeding for 50 s (or increasing reaction times from 1 to 5 min) before adding HpaI in excess and incubating the mixes for at least 1 h at 37 °C in a final reaction volume of 20 µL. Reaction products were analyzed by electrophoresis alongside a 1-kb DNA Ladder in 1% SeaKem LE agarose gels prepared and ran in 1X TAE with ethidium bromide. All the experiments were repeated twice.

### Methylation motif distribution analysis

We obtained the genome assemblies for the two *B. cenocepacia* strains from the NCBI’s Genbank database, under accession numbers AM747720.1 - AM747723.1 (for strain J2315) and Z_CP053300.1 - Z_CP053303.1 (for strain K56-2). The GTWWAC sites were identified by employing the seqkit program^32^ (v.0.15), and their distribution was visualized via a circos plot, which was created using the circlize package in R. In order to identify the homolog of M.BceJIV, we conducted a BLASTP search against the PDB database, using the M.BceJIV amino acid sequence as the query. Expected frequency of the methylation motif was estimated using Markov chain motif distribution analysis as previously described^8^

### Isothermal titration calorimetry assays

The isothermal titration calorimetry experiments were performed on a MicroCal PEAQ-ITC (Malvern Panalytical). The ligands (400 µM) were loaded in the syringe and titrated into 40 µM M.BceJIV**Δ**29 in the cell. Titrations were performed at 25 °C with the standard 10 μcal/s reference power at 600 rpm. The titrations consisted of an initial injection of 0.4 µL (excluded from analysis) followed by 15 injections of 2.5 µL ligand at a rate of 0.5 µL/s at 180 s time intervals. Care was taken to ensure buffer match for the ligand and M.BceJIV**Δ**29 to eliminate heat from buffer mismatch. Data was fit into a single binding site model and analyzed using the MicroCal PEAQ-ITC Analysis software v1.41. All the experiments were repeated in duplicates and the average value was reported.

### Docking of SAM and SAH to the sinefungin’s binding sites in the crystal structure of the M.BceJIV-DNA dimeric complex

The ligands and the protein-DNA dimeric complexes were prepared for docking using the default protocols in LigPrep and the Protein Preparation Wizard from the Schrödinger 2020-4 suite,^33^ respectively. Briefly, a docking grid with edge lengths of 12 Å × 13 Å × 17 Å was generated and centered on the center of mass of sinefungin, using the Schrodinger′s Receptor Grid Generation module^33^. Both SAM and SAH were docked using Glide with the Standard Precision (SP) scoring function^33^ after validating the protocol with sinefungin as a control.

### Energy minimization

Energy minimizations were carried out using GROMACS (version 2021.4)^34^ to assess the stability of the crystallographic binding pose of sinefungin and the predicted binding poses of SAM and SAH in the same binding pocket of protein-DNA complexes embedded in an explicit water environment. Specifically, all three ligand-bound protein-DNA dimeric complexes were embedded in a cubic box with a 10.8 nm edge length in each dimension filled with TIP3P water molecules and 0.15 M NaCl to neutralize the systems, using the *Solution Builder* tool in CHARMM-GUI^35^. The CHARMM36m force field^36^ was used to parameterize the protein, DNA, and solvent in the systems. While the parameters for SAM were present in the July 2022 release of the force field, the parameters for SAH and sinefungin had to be generated using the CHARMM General Force Field (CGenFF) webserver (https://cgenff.silcsbio.com/)^37^. After performing energy minimization cycles for 1,000,000 steps with positional restraints on each of the ligand-bound protein-DNA systems, additional energy minimization cycles in which the systems were unrestrained were conducted for 1,000,000 steps.

### Methylation kinetics assays

The SAM-dependent DNA methylation by full-length M.BceJIV was measured using Promega MTase-Glo^TM^ luminescence assay^38^ where the SAH product is converted to ADP and then ATP. The 20 µL reaction mix contained 20 nM M.BceJIV, increasing concentrations (0 µM to 23 µM) of SAM, and 20 µM DNA duplex (5’-GACGT<u>GTTA**A**C</u>ATGCGCGTC-3’ and 3’-GCTGCACAATTGTACGCGCA-5’) in the same buffer composition described in the methylation-endonuclease assays section. DNA and SAM were preincubated for 5 min before starting the reaction. The SAH calibration curve was generated using 0, 0.125, 0.25 and 0.5 µM SAH (Supplementary Fig. S2). The reactions were performed at room temperature and terminated after 10 min by adding 5 µL of 0.5% trifluoroacetic acid. Five microliters of 6X MTase-Glo^TM^ reagent was added to the mix and incubated for 30 min before adding 30 µL of MTase-Glo^TM^ detection solution and incubating for additional 30 min. Luminescence was measured with a TECAN Infinite 200 Pro Microplate Reader. The data were analyzed using GraphPad Prism v9. All the experiments were repeated in triplicates.

## Data and code availability

The structure factors and coordinate files for crystal structure of the M.BceJI/DNA/sinefungin ternary complex have been deposited in the Protein Data Bank (PDB) under the accession code 8URK.

## Supporting information

PDB Validation Report

## Acknowledgements

This work was funded by grants R35-GM131780 (to A.K.A) and R35GM13965 (to G.F) from the National Institutes of Health (NIH). We thank the staff at the National Synchrotron Light Source II (NSLS-II) beamlines 17-ID-1 and 17-ID-2 for facilitating X-ray data collection. NSLS-II is a United States Department of Energy (DOE) Office of Science User Facility operated for the DOE Office of Science by Brookhaven National Laboratory under Contract No. DE-SC0012704. The Center for BioMolecular Structure (CBMS) at NSLS-II is primarily supported by the NIH, National Institute of General Medical Sciences (NIGMS) through a Center Core P30 Grant (P30GM133893), and by the DOE Office of Biological and Environmental Research (KP1605010). Computational resources needed for this work were provided in part by Scientific Computing at the Icahn School of Medicine at Mount Sinai, the Clinical and Translational Science Awards (CTSA) grant UL1TR004419 from the National Center for Advancing Translational Sciences, and the Office of Research Infrastructure of the National Institutes of Health under award number S10OD026880.

## Author contributions

A.K.A, G.F, M.F and J.K designed the study; R.Q-F performed the crystallographic, biophysical and biochemical studies; J.K assisted and guided the crystallographic, biophysical and biochemical studies; M.N performed the methylation motif distribution analysis; R.G and L.S-E performed and analyzed the energy minimization and docking studies; O.R. assisted in kinetic analysis; G.F guided the methylation motif distribution analysis; M.F. guided the analysis of the energy minimization and docking studies; A.K.A guided the overall project; A.K.A, M.F, G.F, J.K and R.Q-F wrote the manuscript with input from all the authors.

## References

1 Loutet, S. A. & Valvano, M. A. A decade of Burkholderia cenocepacia virulence determinant research. Infect Immun 78, 4088–4100, doi:10.1128/IAI.00212-10 (2010).

2 Roberts, R. J., Vincze, T., Posfai, J. & Macelis, D. REBASE--enzymes and genes for DNA restriction and modification. Nucleic Acids Res 35, D269–270 (2007).

3 Beaulaurier, J., Schadt, E. E. & Fang, G. Deciphering bacterial epigenomes using modern sequencing technologies. Nat Rev Genet 20, 157–172, doi:10.1038/s41576-018-0081-3 (2019).

4 Oliveira, P. H. & Fang, G. Conserved DNA Methyltransferases: A Window into Fundamental Mechanisms of Epigenetic Regulation in Bacteria. Trends Microbiol 29, 28–40, doi:10.1016/j.tim.2020.04.007 (2021).

5 Zhou, J., Horton, J. R., Blumenthal, R. M., Zhang, X. & Cheng, X. Clostridioides difficile specific DNA adenine methyltransferase CamA squeezes and flips adenine out of DNA helix. Nat Commun 12, 3436, doi:10.1038/s41467-021-23693-w (2021).

6 Zhou, J. et al. Repurposing epigenetic inhibitors to target the Clostridioides difficile-specific DNA adenine methyltransferase and sporulation regulator CamA. Epigenetics, 1–12, doi:10.1080/15592294.2021.1976910 (2021).

7 Anton, B. P. & Roberts, R. J. Beyond Restriction Modification: Epigenomic Roles of DNA Methylation in Prokaryotes. Annu Rev Microbiol 75, 129–149, doi:10.1146/annurev-micro-040521-035040 (2021).

8 Oliveira, P. H. et al. Epigenomic characterization of Clostridioides difficile finds a conserved DNA methyltransferase that mediates sporulation and pathogenesis. Nat Microbiol 5, 166–180, doi:10.1038/s41564-019-0613-4 (2020).

9 Zhou, J. et al. Systematic Design of Adenosine Analogs as Inhibitors of a Clostridioides difficile-Specific DNA Adenine Methyltransferase Required for Normal Sporulation and Persistence. J Med Chem 66, 934–950, doi:10.1021/acs.jmedchem.2c01789 (2023).

10 Vandenbussche, I. et al. DNA Methylation Epigenetically Regulates Gene Expression in Burkholderia cenocepacia and Controls Biofilm Formation, Cell Aggregation, and Motility. mSphere 5, doi:10.1128/mSphere.00455-20 (2020).

11 Malone, T., Blumenthal, R. M. & Cheng, X. Structure-guided analysis reveals nine sequence motifs conserved among DNA amino-methyltransferases, and suggests a catalytic mechanism for these enzymes. J Mol Biol 253, 618–632, doi:S0022-2836(85)70577-3 [pii]10.1006/jmbi.1995.0577 (1995).

12 Goedecke, K., Pignot, M., Goody, R. S., Scheidig, A. J. & Weinhold, E. Structure of the N6-adenine DNA methyltransferase M.TaqI in complex with DNA and a cofactor analog. Nat Struct Biol 8, 121–125 (2001).

13. Halford, S. E. et al. Restriction endonuclease reactions requiring two recognition sites. Biochem Soc Trans 27, 696–699, doi:10.1042/bst0270696 (1999).

14 Vanamee, E. S., Santagata, S. & Aggarwal, A. K. FokI requires two specific DNA sites for cleavage. J Mol Biol 309, 69–78 (2001).

15 Vanamee, E. S. et al. A view of consecutive binding events from structures of tetrameric endonuclease SfiI bound to DNA. Embo J 24, 4198–4208 (2005).

16 Shen, B. W. et al. Coordination of phage genome degradation versus host genome protection by a bifunctional restriction-modification enzyme visualized by CryoEM. Structure 29, 521–530 e525, doi:10.1016/j.str.2021.03.012 (2021).

17 Morgan, R. D., Dwinell, E. A., Bhatia, T. K., Lang, E. M. & Luyten, Y. A. The MmeI family: type II restriction-modification enzymes that employ single-strand modification for host protection. Nucleic Acids Res 37, 5208–5221, doi:gkp534 [pii]10.1093/nar/gkp534 (2009).

18 Scavetta, R. D. et al. Structure of RsrI methyltransferase, a member of the N6-adenine beta class of DNA methyltransferases. Nucleic Acids Res 28, 3950–3961 (2000).

19. Osipiuk, J., Walsh, M. A. & Joachimiak, A. Crystal structure of MboIIA methyltransferase. Nucleic Acids Res 31, 5440–5448 (2003).

20 Horowitz, S. et al. Conservation and functional importance of carbon-oxygen hydrogen bonding in AdoMet-dependent methyltransferases. J Am Chem Soc 135, 15536–15548, doi:10.1021/ja407140k (2013).

21 Albanese, K. I. et al. Comparative Analysis of Sulfonium-pi, Ammonium-pi, and Sulfur-pi Interactions and Relevance to SAM-Dependent Methyltransferases. J Am Chem Soc 144, 2535–2545, doi:10.1021/jacs.1c09902 (2022).

22 Urig, S. et al. The Escherichia coli dam DNA methyltransferase modifies DNA in a highly processive reaction. J Mol Biol 319, 1085–1096, doi:10.1016/S0022-2836(02)00371-6 (2002).

23 Woodcock, C. B., Yakubov, A. B. & Reich, N. O. Caulobacter crescentus Cell Cycle-Regulated DNA Methyltransferase Uses a Novel Mechanism for Substrate Recognition. Biochemistry 56, 3913–3922, doi:10.1021/acs.biochem.7b00378 (2017).

24 Horton, J. R. et al. The cell cycle-regulated DNA adenine methyltransferase CcrM opens a bubble at its DNA recognition site. Nat Commun 10, 4600, doi:10.1038/s41467-019-12498-7 (2019).

25 Gupta, Y. K., Chan, S. H., Xu, S. Y. & Aggarwal, A. K. Structural basis of asymmetric DNA methylation and ATP-triggered long-range diffusion by EcoP15I. Nat Commun 6, 7363, doi:ncomms8363 [pii]10.1038/ncomms8363 (2015).

26 Ferreira de Freitas, R., Ivanochko, D. & Schapira, M. Methyltransferase Inhibitors: Competing with, or Exploiting the Bound Cofactor. Molecules 24, doi:10.3390/molecules24244492 (2019).

27 Kaniskan, H. U., Martini, M. L. & Jin, J. Inhibitors of Protein Methyltransferases and Demethylases. Chem Rev 118, 989–1068, doi:10.1021/acs.chemrev.6b00801 (2018).

28 Jumper, J. et al. Highly accurate protein structure prediction with AlphaFold. Nature 596, 583–589, doi:10.1038/s41586-021-03819-2 (2021).

29 Emsley, P. & Cowtan, K. Coot: model-building tools for molecular graphics. Acta Crystallogr D Biol Crystallogr 60, 2126–2132 (2004).

30 Adams, P. D. et al. PHENIX: a comprehensive Python-based system for macromolecular structure solution. Acta Crystallogr D Biol Crystallogr 66, 213–221, doi:S0907444909052925 [pii]10.1107/S0907444909052925 (2010).

31 Moriarty, N. W., Grosse-Kunstleve, R. W. & Adams, P. D. electronic Ligand Builder and Optimization Workbench (eLBOW): a tool for ligand coordinate and restraint generation. Acta Crystallogr D Biol Crystallogr 65, 1074–1080, doi:10.1107/S0907444909029436 (2009).

32 Shen, W., Le, S., Li, Y. & Hu, F. SeqKit: A Cross-Platform and Ultrafast Toolkit for FASTA/Q File Manipulation. PLoS One 11, e0163962, doi:10.1371/journal.pone.0163962 (2016).

33. LigPrep, Protein Preparation Wizard, QikProp, and Glide. (2020).

34 Berendsen, H. J. C., Vanderspoel, D. & Vandrunen, R. Gromacs - a Message-Passing Parallel Molecular-Dynamics Implementation. Comput Phys Commun 91, 43–56, doi:Doi 10.1016/0010-4655(95)00042-E (1995).

35 Jo, S., Kim, T., Iyer, V. G. & Im, W. CHARMM-GUI: a web-based graphical user interface for CHARMM. J Comput Chem 29, 1859–1865, doi:10.1002/jcc.20945 (2008).

36 Huang, J. et al. CHARMM36m: an improved force field for folded and intrinsically disordered proteins. Nat Methods 14, 71–73, doi:10.1038/nmeth.4067 (2017).

37 Vanommeslaeghe, K. et al. CHARMM general force field: A force field for drug-like molecules compatible with the CHARMM all-atom additive biological force fields. J Comput Chem 31, 671–690, doi:10.1002/jcc.21367 (2010).

38 Hsiao, K., Zegzouti, H. & Goueli, S. A. Methyltransferase-Glo: a universal, bioluminescent and homogenous assay for monitoring all classes of methyltransferases. Epigenomics 8, 321–339, doi:10.2217/epi.15.113 (2016).

