## Supplementary figures and images for "*Burkholderia cenocepacia* epigenetic regulator M.BceJIV simultaneously engages two DNA recognition sequences for methylation"

### Supplemental Figure S1

**Supp. Figure 1**

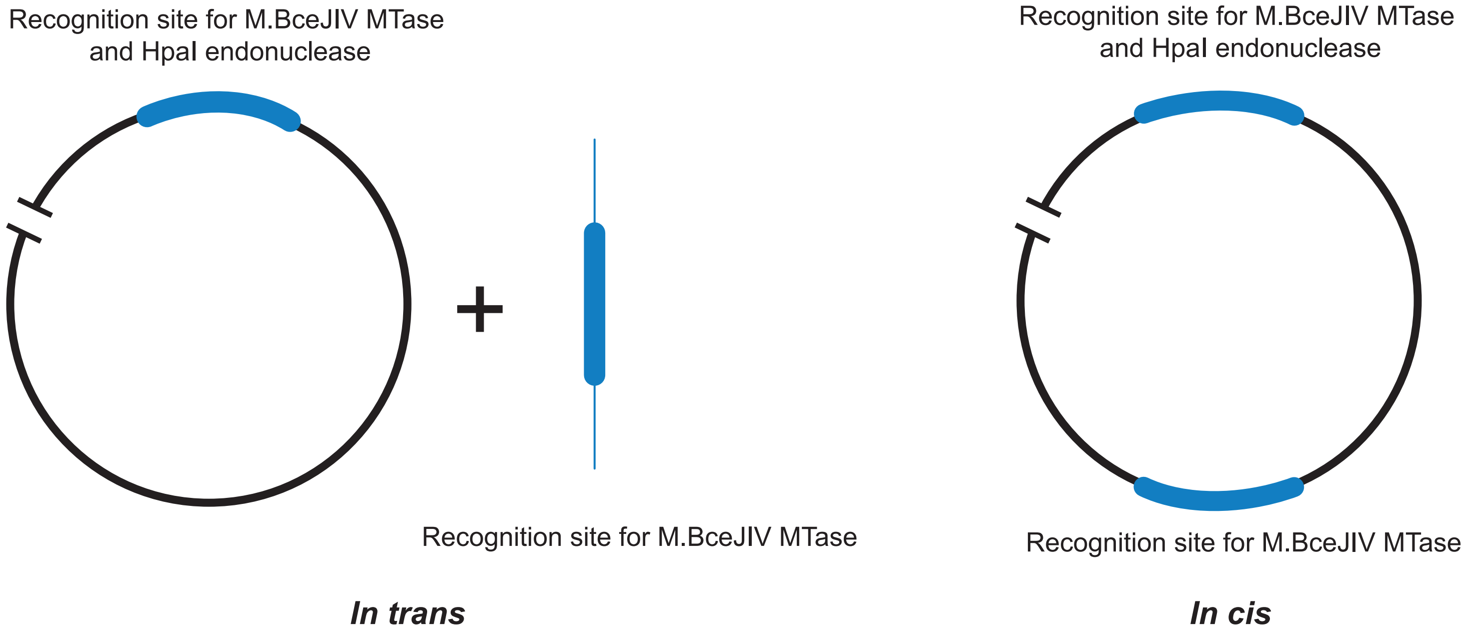

### Supplemental Figure S2

Supp. Figure 2

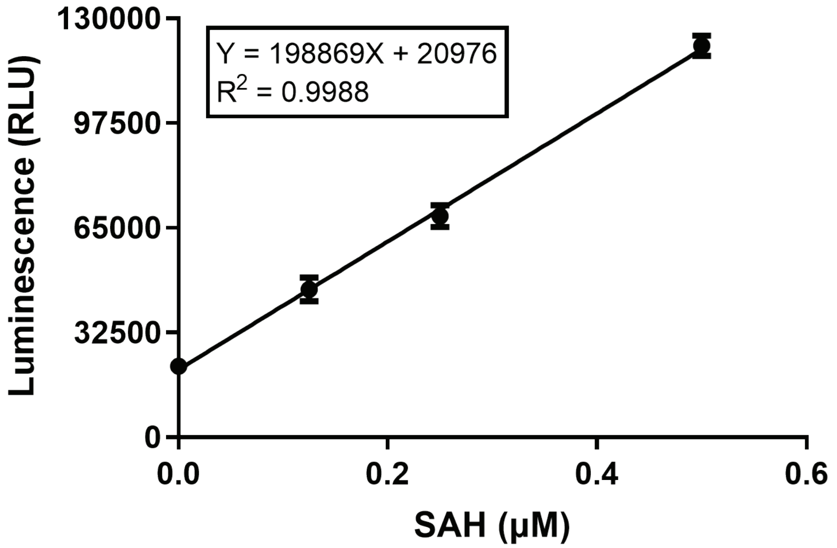
