## Supplementary material for "*Burkholderia cenocepacia* epigenetic regulator M.BceJIV simultaneously engages two DNA recognition sequences for methylation": Table S1

| | M.BceJIV -SFG-DNA complex |
|---|---|
| **Data collection** | |
| Space group | I 41 |
| Cell dimensions | |
| *a*, *b*, *c* (Å) | 137.7 137.7 167.3 |
| α, β, γ (°) | 90.00 90.00 90.00 |
| Resolution (Å) | 97.4-2.11 (2.38-2.11)* |
| $R_{sym}$ or $R_{merge}$ | 10.1 (95.2) |
| $I / \sigma I$ | 9.2 (1.7) |
| Completeness (%) | 94.6 (85.0) |
| Redundancy | 4.7 (4.9) |
| CC(1/2) | 0.99 (0.57) |
| **Refinement** | |
| Resolution (Å) | 44.3-2.11 |
| No. reflections | 49288 |
| $R_{work}$ / $R_{free}$ | 18.8/23.7 |
| No. atoms | |
| Macromolecules | 10195 |
| Ligands | 230 |
| Water | 625 |
| *B*-factors | |
| Macromolecules | 36.24 |
| Ligand/ion | 36.56 |
| Water | 33.45 |
| R.m.s. deviations | |
| Bond lengths (Å) | 0.01 |
| Bond angles (°) | 0.82 |
| Ramachandran Plot | |
| Favored (%) | 92.2 |
| Allowed (%) | 7.28 |
| Outliers (%) | 0.51 |

**Table S2.** Ligand interactions with protein residues, DNA bases, or water molecules within 4 Å from ligand's heavy atoms after energy minimization of the ligand-protein-DNA complexes in an explicit water environment. Interaction types are classified as hydrogen donor (HDonor) or hydrogen acceptor (HAcceptor) between neutral (n) or charged (c) atoms, or they can be salt, hydrophobic, or π-edge interactions.

| **Protein(Chain A)/DNA/Water** | | **Interaction Type** | **Sinefungin** | | | **SAM** | | |
|---|---|---|---|---|---|---|---|---|
| **Residue** | **Atom** | | **Atom** | **Distance (Å)** | **Angle (°)** | **Atom** | **Distance (Å)** | **Angl (°)** |
| Adenine7 | N6 | HDonor nc | HE3 | 2.28 | 96.5 | — | — | — |
| | C6 | Hydrophobic | — | — | — | CE | 3.78 | N/A |
| ASP40 | OD1 | HDonor nn | HN62 | 1.75 | 121.8 | HN62 | 1.71 | 123 |
| PHE41 | HN | HAccep nn | N1 | 2.08 | 116.3 | N1 | 2.02 | 117.4 |
| ASP59 | OD2 | HDonor cc | HN1 | 1.67 | 118 | HN1 | 1.59 | 122.6 |
| | | | HE1 | 2.15 | 90.1 | — | — | — |
| | | Salt | N | 2.71 | N/A | N | 2.63 | N/A |
| | | | NE | 3.02 | N/A | SD | 4.37 | N/A |
| | O | HDonor cc | HE2 | 1.80 | 134.3 | — | — | — |
| | CB | Hydrophobic | — | — | — | — | — | — |
| TYR68 | OH | HDonor nc | HE3 | 2.18 | 101.8 | — | — | — |
| | CE1 | Hydrophobic | — | — | — | SD | 4.0 | N/A |
| ASN70 | HD21 | HAccep nn | O2' | 2.22 | 121.2 | O2' | 2.26 | 123.4 |
| HIS214 | HE2 | HAccep nn | O3' | 1.87 | 118.7 | O3' | 2.03 | 118.2 |
| LYS218 | HN | HAccep nc | OXT | 1.94 | 127.4 | OXT | 1.90 | 120.8 |
| ASP239 | OD2 | Salt | N | 4.864 | N/A | N | 4.782 | N/A |
| PHE241 | CG | PiEdge | N1 | 4.92 | 86.2 | N9 | 5.00 | 84.9 |
| | | | C4 | 5.00 | 88.6 | C5 | 4.87 | 88 |
| | CB | Hydrophobic | C5' | 4.00 | N/A | C5' | 4.00 | N/A |
| SER244 | HN | HAccep nn | O | 1.86 | 125.1 | O | 1.89 | 127.2 |
| | HG1 | HAccep nn | O | 1.88 | 160 | O | 1.83 | 164 |
| THR246 | HG1 | HAccep nc | OXT | 1.77 | 121.4 | OXT | 1.70 | 115.2 |
| GLU262 | OE1 | HDonor nn | HO3' | 1.67 | 122.4 | HO3' | 1.66 | 118.6 |
| | OE2 | HDonor cn | HO2' | 1.71 | 114.9 | HO2' | 1.69 | 116.8 |
| ILE263 | HN | HAccep nn | N3 | 2.66 | 100.1 | N3 | 2.66 | 102.8 |
| Water1 | H1 | HAccep nn | N7 | 1.90 | 112.4 | — | — | — |
| Water2 | OH2 | HDonor nc | HN3 | 1.74 | 123.5 | HN3 | 1.73 | 125.4 |
